# Prenatal 3i-0777 treatment attenuates left ventricular hypertrabeculation and dysfunction in a retinoic acid–induced mouse model

**DOI:** 10.64898/2026.09.03.748426

**Authors:** Qinglan Shu, Zhengyang Wang, Qingguo Meng, Xinyi Liu, Lixue Yin

**Author notes:** Correspondence: Lixue Yin.

## Abstract

Excessive left ventricular trabeculation and thinning of the compacted myocardium arise during cardiac development and may be accompanied by myocardial dysfunction. Gestational exposure to all-trans retinoic acid (ATRA) induces a comparable phenotype in mice, providing a model for evaluating prenatal pharmacological intervention. We examined whether 3i-0777, a small-molecule enhancer of GATA4– NKX2-5 transcriptional synergy, could attenuate these structural and functional abnormalities. Pregnant C57BL/6NCrl dams were randomly assigned to untreated control, vehicle control, ATRA, or ATRA plus 3i-0777 groups (12 independent litters per group). Both ATRA-exposed groups received ATRA (50 mg/kg) on embryonic day (E)12.5. In the ATRA plus 3i-0777 group, 3i-0777 (15 mg/kg) was administered as a separate injection concurrently with ATRA at E12.5, followed by 5 mg/kg every 8 h (three doses per day) on E13.5–E15.5. At postnatal day 1, one pup was randomly selected from each litter for echocardiography and subsequent histomorphometry. The echocardiographic and histomorphometric ratios of non-compacted to compacted myocardial layer thickness (N/C ratios) were higher, and the absolute magnitude of global longitudinal strain (|GLS|) was lower, in the ATRA group than in either control group (all adjusted *P* < 0.001). Compared with the ATRA group, the ATRA plus 3i-0777 group had lower echocardiographic and histomorphometric N/C ratios (0.58 ± 0.19 vs 2.19 ± 0.18 and 0.90 ± 0.33 vs 2.46 ± 0.48, respectively), higher |GLS| (25.9 ± 2.5% vs 15.3 ± 2.7%), and a smaller left ventricular internal diameter at end-diastole (1.15 ± 0.06 vs 1.42 ± 0.10 mm; all adjusted *P* < 0.001). These findings provide proof of concept that ATRA-induced left ventricular hypertrabeculation and dysfunction can be pharmacologically attenuated in utero.

## Introduction

Excessive left ventricular trabeculation is a morphological phenotype that may occur as a normal variant or in association with developmental cardiac abnormalities and myocardial dysfunction.^1^ Gestational exposure to all-trans retinoic acid (ATRA) induces ventricular hypertrabeculation and thinning of the compact myocardium in mice,^2,3^ providing an experimental model of abnormal ventricular wall development. We therefore examined whether 3i-0777, a small-molecule enhancer of GATA4–NKX2-5 transcriptional synergy,^4^ could attenuate left ventricular hypertrabeculation and dysfunction at postnatal day 1.

## Methods

C57BL/6NCrl mice were obtained from Beijing Vital River Laboratory Animal Technology Co., Ltd. (Beijing, China). Female and male mice were 10–12 weeks old at mating. The day of vaginal-plug detection was designated embryonic day 0.5 (E0.5). Dams were randomly assigned to one of four groups: untreated control, vehicle control, ATRA, or ATRA plus 3i-0777 (12 independent litters per group).

All-trans retinoic acid (ATRA; catalogue no. R2625) and dimethyl sulfoxide (DMSO; catalogue no. D2438) were purchased from Sigma-Aldrich (St Louis, MO, USA). Corn oil (catalogue no. C805618) was purchased from Shanghai Macklin Biochemical Co., Ltd. (Shanghai, China). 3i-0777 [N-(4-chlorophenyl)-5-methyl-N-(4-methyl-4,5-dihydrothiazol-2-yl)-3-phenylisoxazole-4-carboxamide] (purity ≥90%) was purchased from Enamine Ltd. (Kyiv, Ukraine). ATRA and 3i-0777 were formulated separately in 5% DMSO and 95% corn oil. The vehicle consisted of the same proportions of DMSO and corn oil without active compound.

All injections were administered intraperitoneally at a volume of 8 mL/kg. Both ATRA-exposed groups received ATRA (50 mg/kg) at E12.5. In the ATRA plus 3i-0777 group, an initial dose of 3i-0777 (15 mg/kg) was administered concurrently with ATRA in a separate injection at E12.5, followed by 5 mg/kg every 8 h (three doses per day) on E13.5–E15.5. No additional injections were administered to the ATRA group after E12.5. At E12.5, vehicle control dams received two separate vehicle injections to match the ATRA and initial 3i-0777 injections; thereafter, vehicle was administered every 8 h (three injections per day) on E13.5–E15.5. Untreated controls received no injections.

Treatment was allocated at the dam level, with the litter serving as the experimental unit. At postnatal day 1 (P1), one pup was randomly selected from each litter for analysis. Pup sex was not recorded because it could not be reliably determined by external examination at P1.

At P1, pups were weighed. Echocardiography was performed without anaesthesia using a Vevo 3100 system equipped with an MX550D 40-MHz transducer (FUJIFILM VisualSonics, Toronto, Canada). Images were analysed using Vevo LAB software, version 5.8.2 (FUJIFILM VisualSonics). Left ventricular internal diameter at end-diastole (LVIDd) was measured in parasternal long-axis B-mode images. Left ventricular ejection fraction (EF), fractional shortening (FS), end-diastolic volume (EDV), end-systolic volume (ESV), stroke volume (SV), and global longitudinal strain (GLS) were derived from parasternal long-axis B-mode cine loops using semi-automated endocardial speckle tracking with Vevo Strain in Vevo LAB. GLS was reported as an absolute magnitude (|GLS|) and averaged over three consecutive cardiac cycles. At end-systole, the thicknesses of the non-compacted (N) and compacted (C) myocardial layers were measured in left ventricular short-axis images at the papillary-muscle level. The N/C ratio was calculated as non-compacted myocardial layer thickness divided by compacted myocardial layer thickness.

After echocardiography, pups were euthanized by rapid decapitation with sharp surgical scissors. Death was confirmed before the hearts were excised and fixed in 4% paraformaldehyde. Hearts were embedded in paraffin, cut into 5-μm sections, and stained with haematoxylin and eosin. Histological sections were digitised using an Axio Scan 7 slide scanner equipped with a 20× objective and analysed in ZEN software, version 3.9 (Carl Zeiss Microscopy GmbH, Oberkochen, Germany). For histomorphometry, the thicknesses of the non-compacted and compacted myocardial layers were measured in short-axis sections at the papillary-muscle level, and the N/C ratio was calculated in the same manner. All echocardiographic and histomorphometric measurements were performed by an observer blinded to group allocation.

Outcomes included body weight; echocardiographic measures (LVIDd, EF, FS, EDV, ESV, SV, |GLS|, non-compacted and compacted myocardial layer thicknesses, and the N/C ratio); and histomorphometric measurements of both myocardial layer thicknesses and the N/C ratio. Data are reported as mean ± standard deviation (SD). Each outcome was analysed using Welch’s one-way analysis of variance, followed by two-sided Games–Howell tests for all pairwise comparisons. Reported pairwise *P* values were adjusted within each outcome using the Games–Howell procedure. A *P* value < 0.05 was considered statistically significant. Statistical analyses were performed using GraphPad Prism version 10.6.1 (GraphPad Software).

## Results

One randomly selected P1 pup from each of the 48 litters (*n* = 12 per group) was included in the analysis; no outcome data were missing. Compared with the untreated and vehicle control groups, the ATRA group had higher echocardiographic N/C ratios (2.19 ± 0.18 vs 0.40 ± 0.07 and 0.35 ± 0.13, respectively), higher histomorphometric N/C ratios (2.46 ± 0.48 vs 0.62 ± 0.14 and 0.59 ± 0.07, respectively), lower | GLS| (15.3 ± 2.7% vs 30.7 ± 6.7% and 27.6 ± 4.3%, respectively), and a larger LVIDd (1.42 ± 0.10 vs 1.10 ± 0.07 and 1.12 ± 0.08 mm, respectively; all *P* < 0.001; Figure 1F and Table 1). Pairwise comparisons between the untreated and vehicle control groups were not statistically significant for these measures.

**Table 1.**
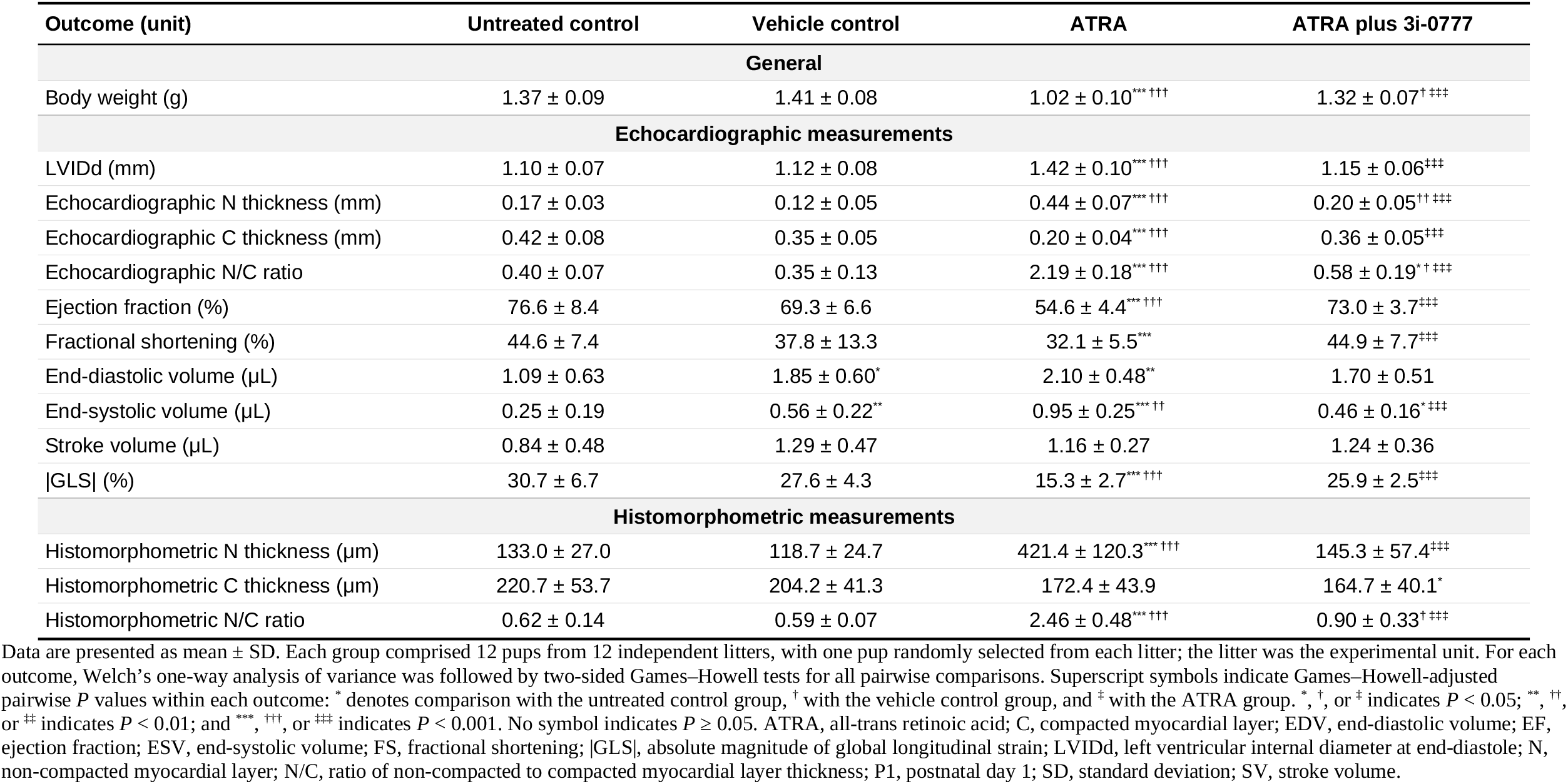
Body weight and cardiac structural and functional outcomes at P1.

**Figure 1.**
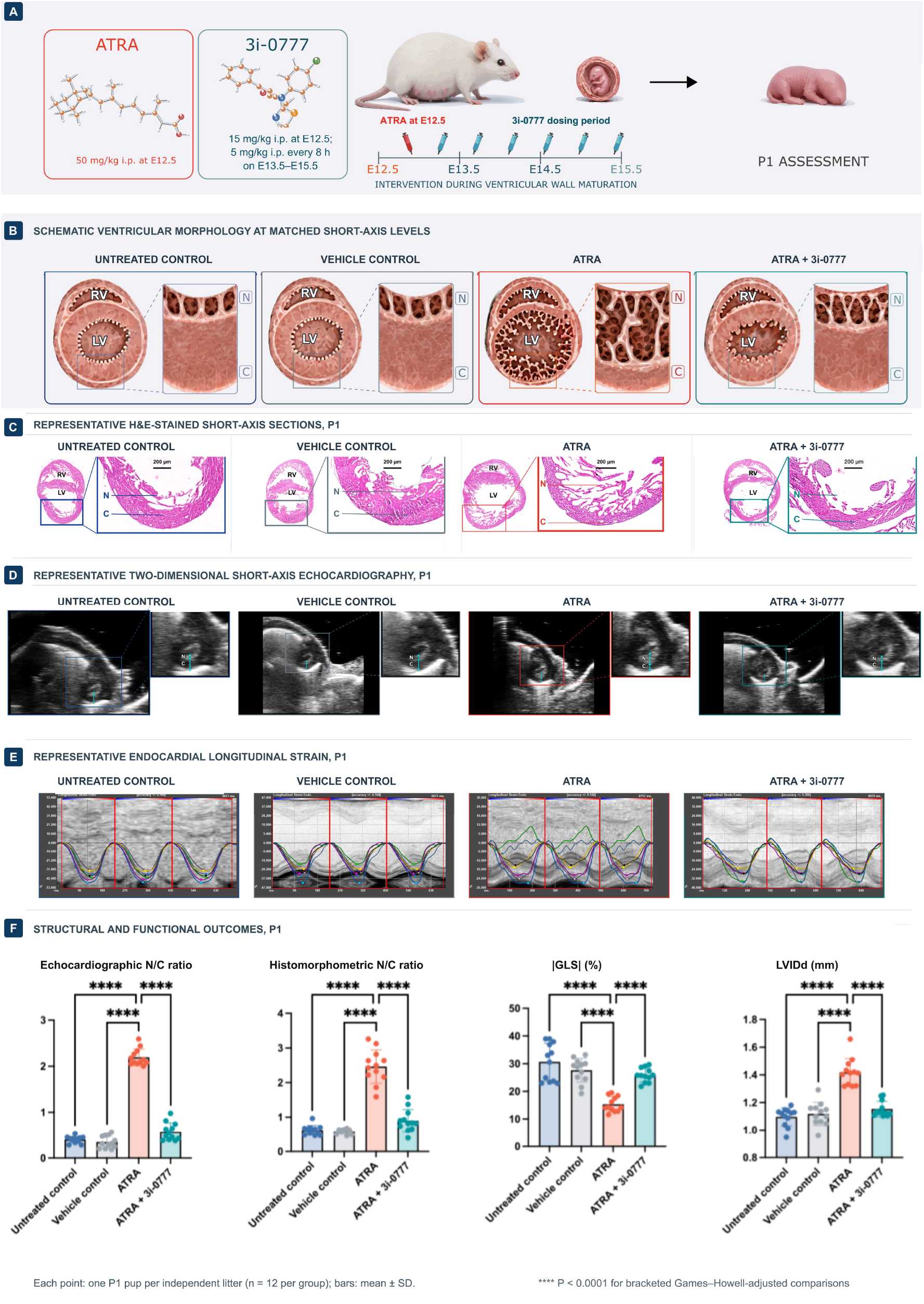
Prenatal 3i-0777 attenuates ATRA-induced left ventricular hypertrabeculation and dysfunction at P1. **(A)** Schematic study design and dosing schedule. Pregnant C57BL/6NCrl dams were randomly assigned to untreated control, vehicle control, ATRA, or ATRA plus 3i-0777 groups (*n* = 12 independent litters per group). Both ATRA-exposed groups received ATRA (50 mg/kg) at E12.5. In the ATRA plus 3i-0777 group, an initial dose of 3i-0777 (15 mg/kg) was administered concurrently with ATRA in a separate injection at E12.5, followed by 5 mg/kg every 8 h (three doses per day) on E13.5–E15.5. No additional injections were administered to the ATRA group after E12.5. At E12.5, vehicle controls received two separate vehicle injections to match the ATRA and initial 3i-0777 injections; thereafter, vehicle was administered every 8 h (three injections per day) on E13.5–E15.5. Untreated controls received no injections. All injections were administered intraperitoneally at a volume of 8 mL/kg. Syringe icons denote dosing periods rather than individual injections. **(B)** Schematic short-axis views illustrating ventricular morphology and the non-compacted and compacted myocardial layers. **(C)** Representative short-axis sections stained with haematoxylin and eosin at the papillary-muscle level. Boxed left ventricular regions are shown at higher magnification. Scale bars in the magnified images, 200 μm. **(D)** Representative two-dimensional short-axis echocardiograms at the papillary-muscle level. Cyan calipers indicate end-systolic thickness measurements of the non-compacted and compacted myocardial layers. **(E)** Representative endocardial longitudinal-strain traces derived from parasternal long-axis cine loops over three consecutive cardiac cycles using Vevo LAB, version 5.8.2. Segmental curves are colour-coded as follows: anterior base, dark blue; anterior mid, yellow; anterior apex, magenta; posterior apex, light blue; posterior mid, white; and posterior base, green. The black curve represents the mean of the six segmental curves, and red vertical lines denote cardiac-cycle boundaries. Negative strain indicates myocardial shortening. **(F)** Echocardiographic N/C ratio, histomorphometric N/C ratio, absolute magnitude of global longitudinal strain (|GLS|), and left ventricular internal diameter at end-diastole (LVIDd). Each point represents one randomly selected P1 pup from an independent litter (*n* = 12 per group); the litter was the experimental unit. Bars show mean ± SD. Group differences were assessed using Welch’s one-way analysis of variance, followed by two-sided Games–Howell tests for all pairwise comparisons. \*\*\*\**P* < 0.0001 for the bracketed comparisons. Pairwise *P* values were Games–Howell adjusted within each outcome. ATRA, all-trans retinoic acid; C, compacted myocardial layer; E, embryonic day; |GLS|, absolute magnitude of global longitudinal strain; LV, left ventricle; N, non-compacted myocardial layer; N/C, ratio of non-compacted to compacted myocardial layer thickness; P1, postnatal day 1; RV, right ventricle; SD, standard deviation.

Prenatal treatment with 3i-0777 attenuated the ATRA-induced cardiac phenotype. Compared with the ATRA group, the ATRA plus 3i-0777 group had lower echocardiographic and histomorphometric N/C ratios (0.58 ± 0.19 vs 2.19 ± 0.18 and 0.90 ± 0.33 vs 2.46 ± 0.48, respectively) and higher |GLS| (25.9 ± 2.5% vs 15.3 ± 2.7%; all *P* < 0.001). The echocardiographic N/C ratio remained higher than in both control groups (both *P* < 0.05), whereas the histomorphometric N/C ratio remained higher than in the vehicle control group (*P* < 0.05); its comparison with the untreated control group was not statistically significant. Pairwise comparisons of |GLS| between the ATRA plus 3i-0777 group and the untreated and vehicle control groups were not statistically significant (*P* = 0.144 and 0.629, respectively; Figure 1F and Table 1).

The ATRA plus 3i-0777 group also had a smaller LVIDd than the ATRA group (1.15 ± 0.06 vs 1.42 ± 0.10 mm; *P* < 0.001). On echocardiography, the ATRA plus 3i-0777 group had a thinner non-compacted myocardial layer (0.20 ± 0.05 vs 0.44 ± 0.07 mm) and a thicker compacted myocardial layer (0.36 ± 0.05 vs 0.20 ± 0.04 mm; both *P* < 0.001). The non-compacted layer remained thicker than in the vehicle control group (*P* < 0.01). Histomorphometry likewise showed a thinner non-compacted myocardial layer in the ATRA plus 3i-0777 group than in the ATRA group (145.3 ± 57.4 vs 421.4 ± 120.3 μm; *P* < 0.001). Histomorphometric compacted myocardial layer thickness did not differ between the ATRA plus 3i-0777 and ATRA groups (164.7 ± 40.1 vs 172.4 ± 43.9 μm; *P* = 0.969), and was lower in the ATRA plus 3i-0777 group than in the untreated control group (164.7 ± 40.1 vs 220.7 ± 53.7 μm; *P* < 0.05; Table 1).

EF and FS were higher in the ATRA plus 3i-0777 group than in the ATRA group (73.0 ± 3.7% vs 54.6 ± 4.4% and 44.9 ± 7.7% vs 32.1 ± 5.5%, respectively; both *P* < 0.001). Pairwise comparisons of EF and FS between the ATRA plus 3i-0777 group and either control group were not statistically significant. ESV was lower in the ATRA plus 3i-0777 group than in the ATRA group (0.46 ± 0.16 vs 0.95 ± 0.25 μL; *P* < 0.001), although it remained higher than in the untreated control group (*P* < 0.05). EDV was higher in the vehicle and ATRA groups than in the untreated control group (*P* < 0.05 and *P* < 0.01, respectively), but the comparison between the ATRA plus 3i-0777 and ATRA groups was not statistically significant. No pairwise comparisons in SV were statistically significant. Body weight was higher in the ATRA plus 3i-0777 group than in the ATRA group (1.32 ± 0.07 vs 1.02 ± 0.10 g; *P* < 0.001), but remained lower than in the vehicle control group (*P* < 0.05); its comparison with the untreated control group was not statistically significant (Table 1).

## Discussion

Prenatal 3i-0777 treatment during ventricular wall maturation attenuated the ATRA-induced cardiac phenotype in P1 pups. Treatment lowered echocardiographic and histomorphometric N/C ratios, reduced non-compacted myocardial layer thickness by both methods, and decreased LVIDd. Systolic function also improved, with higher |GLS|, EF, and FS and a lower ESV. The concordant reductions in N/C ratio and non-compacted layer thickness, together with improvements in deformation-based and conventional indices, support attenuation of both left ventricular hypertrabeculation and associated dysfunction. These data provide proof of concept that an ATRA-induced developmental cardiac phenotype can be pharmacologically modified in utero.

Ventricular wall maturation depends on expansion of the compact myocardium, trabecular remodelling, cardiomyocyte proliferation, and coordinated endocardial–myocardial signalling.^5,6^ During early cardiogenesis, excess retinoic acid signalling can disrupt differentiation of cardiac progenitors toward ventricular cardiomyocytes, dysregulate fibroblast growth factor signalling, and impair cell-cycle exit.^7^ Previous studies identified 3i-0777 as an enhancer of GATA4–NKX2-5 transcriptional synergy in reporter assays^4^ and showed that it augments stretch-induced atrial natriuretic peptide expression in neonatal rat cardiomyocytes.^8^ *Gata4* and *Nkx2-5* have established roles in ventricular development.^9,10^ The present findings are consistent with this developmental biology and support further investigation of the GATA4–NKX2-5 transcriptional network in the cardiac response to 3i-0777.

## Competing interests

Qinglan Shu and Lixue Yin are named inventors on Chinese patent application CN202610233138.9 (publication CN121796393A), assigned to Sichuan Academy of Medical Sciences and Sichuan Provincial People’s Hospital, relating to the use of 3i-0777 for the prevention and/or treatment of left ventricular non-compaction. All other authors declare no competing interests.

## Data availability

The data supporting the findings of this study are available from the corresponding author upon reasonable request.

## Funding

This work was supported by the National Natural Science Foundation of China (grant number 82472005).

## Ethical approval

Animal procedures were approved by the Ethics Committee for Basic and Clinical Research of Sichuan Academy of Medical Sciences and Sichuan Provincial People’s Hospital (approval no. 2026-150).

